# Activation of NF-κB signaling by optogenetic clustering of IKKα and β

**DOI:** 10.1101/2024.06.12.598631

**Authors:** Alexandra A.M. Fischer, Merlin M. Grimm, Manfred Fliegauf, Bodo Grimbacher, Sven Rahmann, Wilfried Weber

## Abstract

A large percentage of proteins form higher-order structures in order to fulfill their function. These structures are crucial for the precise spatial and temporal regulation of the cellular signaling network. Investigation of this network requires sophisticated research tools, such as optogenetic tools, that allow dynamic control over the signaling molecules. Cryptochrome 2 and its variations are the best-characterized oligomerizing photoreceptors the optogenetics toolbox has to offer. Therefore, we utilized this switch and combined it with an eGFP-binding nanobody, to build a toolbox of optogenetic constructs that enables the oligomerization of any eGFP-tagged protein of interest. We further introduced the higher clustering variant Cry2_olig_ and an intrinsically disordered region to create higher-order oligomers or phase-separated assemblies to investigate the impact of different oligomerization states on eGFP-tagged signaling molecules. We apply these constructs to cluster IKKα and IKKβ, which resemble the central signaling integrator of the NF-κB pathway, thereby engineer a potent, blue-light-inducible activator of NF-κB signaling.

## Introduction

Protein oligomerization is an essential mechanism for the control of cellular signaling processes and a large percentage of cellular proteins oligomerize in order to fulfill their function^1–3^. Their correct assembly, in terms of location, time, order of oligomerization, and binding partners, decides critically over the signaling output^4^. Investigation of these processes requires sophisticated research tools capable of tuning oligomerization. A versatile method to achieve this is optogenetics. This method uses plant-derived photoreceptors, genetically fused to cellular proteins and thereby enables control over their function using light of a specific wavelength as a non-invasive, reversible, and dose-dependent stimulus. The optogenetics toolbox provides a large variety of photoswitches that allow the assembly of a protein of interest into homo-or heterodimers^5–9^, homo- or heterooligomers^10–12^, or phase-separated biomolecular condensates^13–15^. The most established oligomerizing photoreceptor is Cryptochrome 2 (Cry2)^10^. It can form heterodimers with its binding partner CIBN^5^ and furthermore, variants that can form higher-order oligomers (Cry2_olig_^16^ or Cry2_clust_^17^) have been engineered.

Combining Cry2 with an intrinsically disordered region (IDR) such as the N-terminal domain of Fused in Sarcoma (FUS_N_) leads to the formation of phase-separated liquid-like biomolecular condensates, called optoDroplets, after blue light illumination. The exchange of Cry2 with the Cry2_olig_ variant leads to the formation of larger, solid-like gels^14^.

Cry2, fused to different signaling molecules, has been used in many studies to dynamically control signaling pathways^18,19^, such as the mitogen-activated protein kinase (MAPK)^20–23^ pathway, Wnt/β-catenin signaling^10,24^, receptor tyrosine kinase (RTK)^25–27^ activated signaling pathways, or recently as well NF-κB signaling by clustering MyD88 and TRAF6^28^.

NF-κB signaling is regulated by higher-order oligomerization events at several levels of the pathway^29^. It is activated by various factors, among them TNF-α which binds to the tumor necrosis factor receptor (TNFR). Stimulation of the TNFR with TNF-α leads to receptor trimerization and the recruitment of a higher-order complex consisting, among others, of adaptor proteins serine/threonine-kinases and E2/E3 ubiquitin ligases, that catalyze the formation of polyubiquitin chains. The TAK1/TAB2/TAB1 complex is recruited to the polyubiquitin chains, as well as the IκB kinase (IKK) complex. The IKK complex represents the central signal integration module of the NF-κB signaling pathway and consists of the regulatory subunit NEMO, which acts as a scaffold to assemble the two serine-threonine kinases IKKα and IKKβ. Recently, it has been discovered that the binding of the polyubiquitin chains (polyUb) to NEMO leads to the formation of phase-separated biomolecular condensates that are crucial for efficient downstream signaling activation. They harbor TAK1 as well as IKKα and β, thereby likely facilitating efficient IKK phosphorylation and activation by TAK1^30,31^. Another widely discussed IKK activation mechanism is trans-autophosphorylation of IKKα and β. Subsequently, IKKβ phosphorylates IκBα, which is then ubiquitinated and degraded. This releases the NF-kB transcription factor dimer (e.g. RelA and p50), which translocates to the nucleus and activates gene expression^32–35^.

Usually, optogenetic tools require the fusion of a photoreceptor to a protein of interest which requires recloning for every new target protein. In this study, we engineered a toolbox of constructs based on the blue-light oligomerizing photoreceptor Cry2 and the recruitment of proteins of interest via an eGFP binding nanobody (NbGFP)^36^. We combine NbGFP with Cry2, Cry2_olig_^16^, or the phase separation inducing optoDroplet configuration (Cry2/Cry2_olig_-FUS_N_)^14^. This enabled the clustering of eGFP-tagged proteins of

interest to different degrees. We applied the constructs to cluster IKKα and β, as their role as central signaling integrators makes them the perfect target for optogenetic regulation of NF-κB signaling. Using these constructs, we were able to uncouple the pathway activation from upstream signaling events and engineer a blue-light inducible, optogenetic activator of the NF-κB pathway.

## Results

### Design of a construct toolbox to gradually cluster eGFP-tagged proteins

We envisioned a toolbox of clustering constructs that enables convenient targeting and clustering of any protein of interest to different degrees. Therefore, we fused the photoreceptor Cry2 to mCherry (mCh) for visualization, a nanobody that can specifically bind to the fluorescent protein eGFP (NbGFP) and a nuclear export sequence (NES) to direct the fusion protein to the cytoplasm (Cry2-mCh-NES-NbGFP). The protein of interest is fused to eGFP, which is a usually well-tolerated fusion partner^37–39^. From this basic design, we went on to mutate Cry2 to Cry2_olig_ to increase clustering (Cry2_olig_-mCh-NES-NbGFP) and in the next step, inspired by the optoDroplet tool, incorporated FUS_N_. We hypothesized that this would lead to the formation of phase-separated bio-molecular condensates after blue-light illumination (Cry2-mCh-FUS_N_-NES-NbGFP and Cry2_olig_-mCh-FUS_N_-NES-NbGFP, Figure 1). For this study, we selected IKKα and IKKβ as target proteins to test our system and tagged them with eGFP. To validate expression and localization of the system components, we transfected them into HEK-293T cells, either eGFP-IKKα and eGFP-IKKβ only or together with one of the clustering constructs, and kept them either in darkness or under blue light. Both, eGFP-IKKα and eGFP-IKKβ, as well as the clustering constructs, localized in the cytoplasm, and all constructs, showed a diffuse signal in darkness. Upon blue light illumination, we observed a low number of small clusters for Cry2-mCh-NES-NbGFP and large cytoplasmic clusters for Cry2_olig_-mCh-NES-NbGFP, Cry2-mCh-FUS_N_-NES-NbGFP and Cry2_olig_-mCh-FUS_N_-NES-NbGFP in both, mCherry and eGFP channel, demonstrating the binding and recruitment of IKKα/β into the clusters (Figure 1).

**Figure 1:**
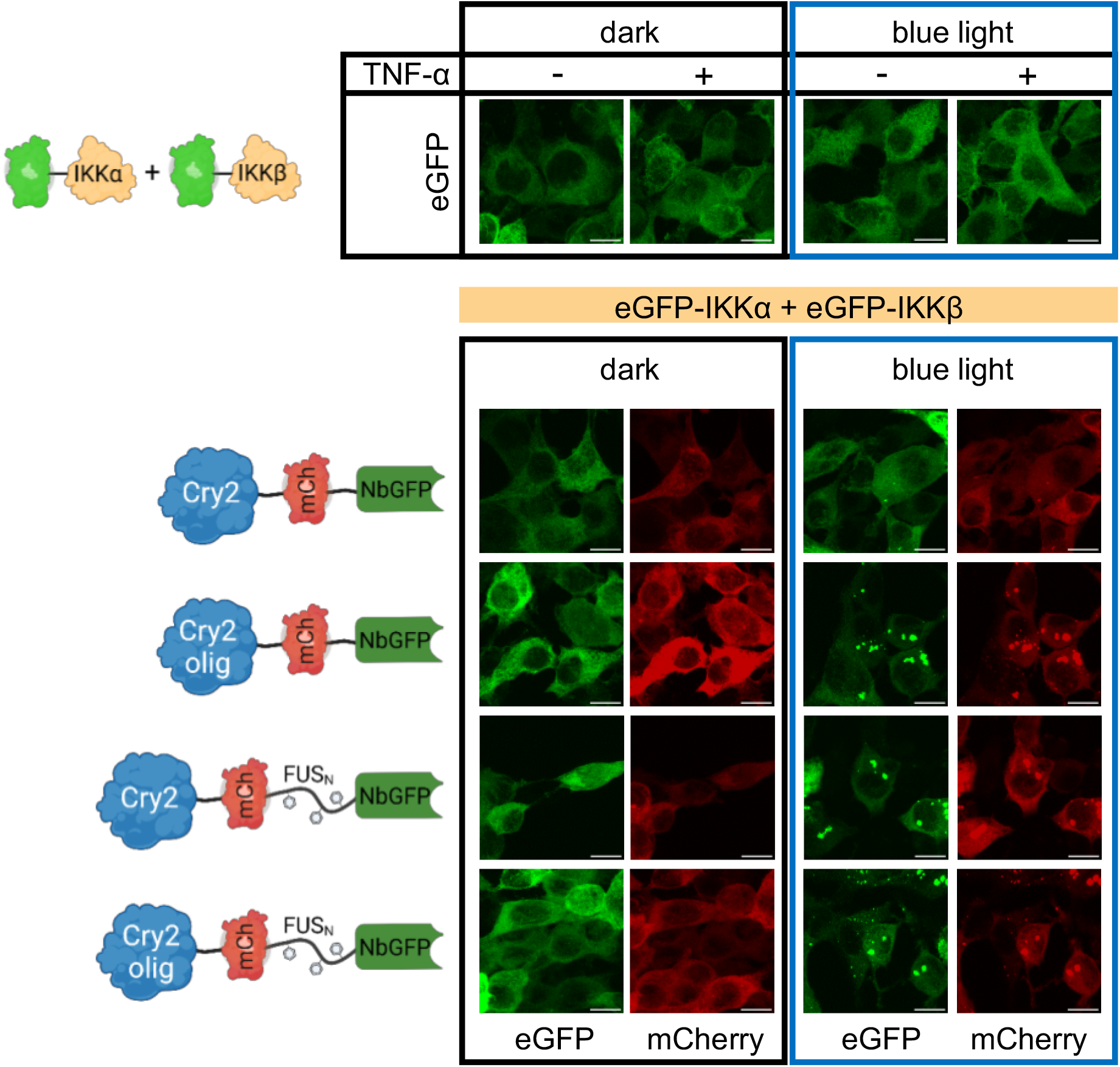
Microscopical characterization of the clustering constructs. HEK-293T cells were transfected with the indicated constructs and an NF-κB-responsive firefly luciferase reporter. Blue-light illumination (5 μmol m^-2^ s^-1^) was started 8 h after transfection and samples were fixed for microscopy analysis 24 h later. Representative images are shown; scale bar = 10 μm.

### Optogenetic clustering of IKKα and IKKβ activates NF-kB signaling independent of upstream signals

We hypothesized that bringing IKKα and β optogenetically into close proximity might be sufficient to activate the kinases independent from upstream activators (Figure 2A). We tested this hypothesis by transfecting HEK-293T cells with eGFP-IKKα and eGFP-IKKβ and together with each construct of the toolbox and an NF-κB-responsive firefly luciferase reporter^40^. Indeed, blue-light illumination led to strong activation of reporter expression. Although we could not reach the total NF-κB reporter signal strength in comparison to TNF-α stimulation, we could achieve similar fold induction, as co-expression of the optogenetic constructs decreased the dark background. This was likely due to the binding of NbGFP to eGFP-IKKα and β (Figure 2B, Supplementary Figure 1). Interestingly, the activation strength depended largely on the clustering construct, and the FUS_N_-containing constructs did not lead to the highest signal output or fold induction. Instead, the Cry2_olig_-mCh-NES-NbGFP construct, which leads to high-order oligomerization, resulted in the highest total signal and highest fold induction of NF-κB activity (Figure 2B).

**Figure 2:**
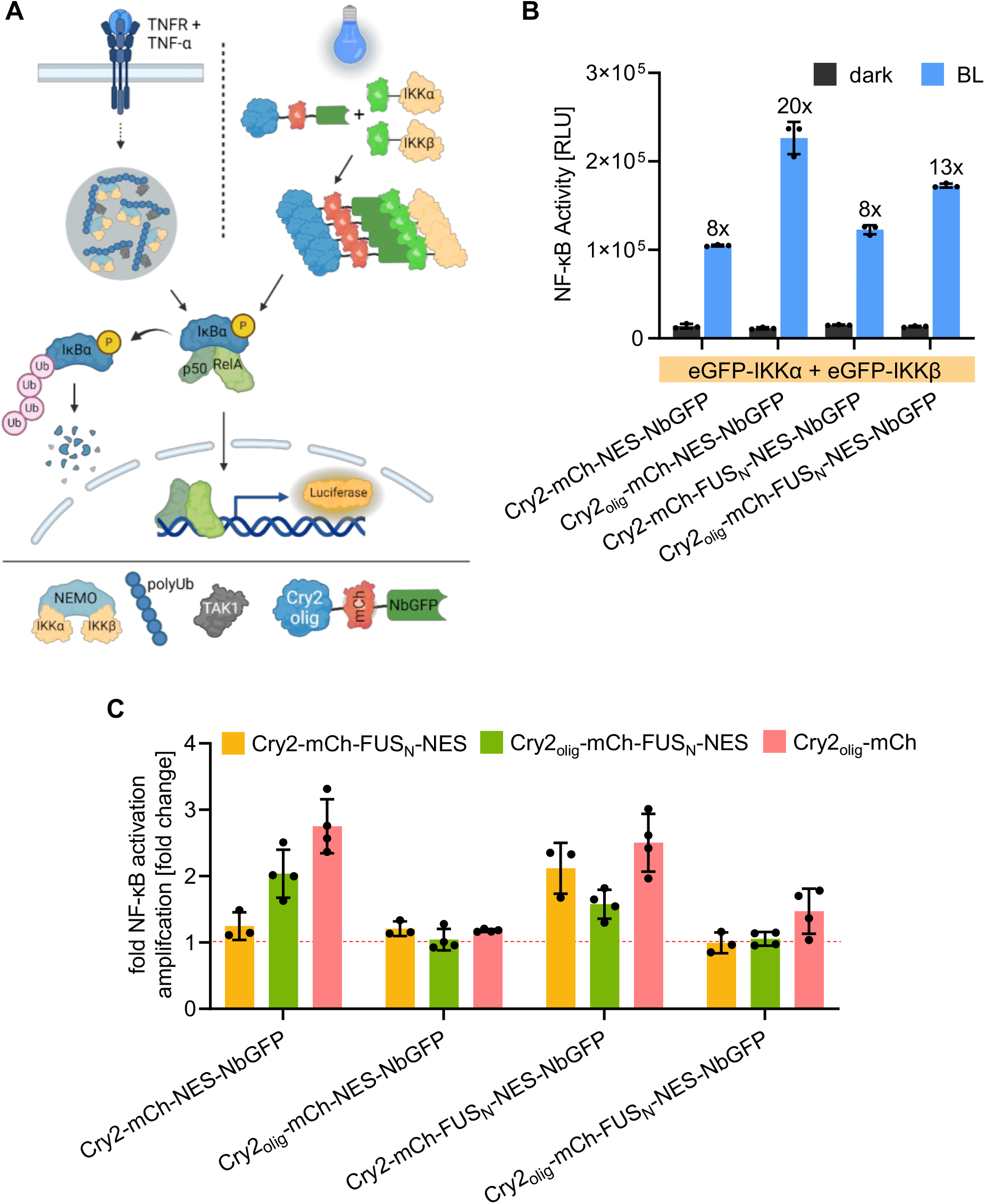
Optogenetic activation of NF-κB signaling. (**A**) Schematic depiction of endogenous and optogenetic NF-κB pathway activation. The TNFR is activated upon binding of TNF-α. After intermediate steps (indicated by the dashed line), binding of NEMO to polyUb chains leads to the formation of biomolecular condensates that also recruit the kinase TAK1 and the IKK complex through binding to the polyUbs or NEMO respectively. The IKK complex is activated and subsequently phosphorylates IκBα, which is then ubiquitinated and degraded. This releases the NF-κB transcription factor dimer RelA/p50, which translocates to the nucleus and activates gene expression. In the optogenetic approach, we cluster eGFP-tagged IKKα and IKKβ and thereby activate the pathway independent from upstream processes. (**B**) Blue-light induced activation of NF-κB signaling. HEK-293T cells were transfected with the indicated constructs and an NF-κB-responsive firefly luciferase reporter. Blue-light illumination (5 μmol m^-2^ s^-1^) was started 8 h after transfection and firefly luciferase activity was determined 24 h later. Mean ± SD of a representative assay is shown (N = 3). Fold NF-κB activation, calculated between the respective dark and BL samples is shown above each BL bar. (**C**) Influence of an additional clustering construct (Cry2-mCh-FUS_N_-NES, Cry2_olig_-mCh-FUS_N_-NES, and Cry2_olig_-mCh) on fold pathway activation. HEK-293T cells were treated as described in B, except that the respective colorindicated constructs were additionally transfected. Firefly activity was measured and fold NF-κB activation was calculated between dark and BL samples. Then the fold signal amplification between samples with or without the clustering-increasing construct was calculated. Single data points represent the fold signal amplification of 3-4 independent experiments. Means ± SD are plotted.

The IKK complex is reported to be recruited into NEMO/polyUbi phase-separated bio-molecular condensates^30,31^. Hence, we wondered, if the introduction of an additional phase-separating construct could increase pathway activation further. To this aim, we designed three additional constructs: first, Cry2-mCh-FUS_N_-NES, second, Cry2_olig_-mCh-FUS_N_-NES and third, Cry2_olig_-mCh. To enable visualization of the colocalization of eGFP-IKKα/β with the mCherry-tagged clustering constructs and the constructs to improve clustering (Cry2_olig_-mCh, Cry2/Cry2_olig_-mCh-FUS_N_-NES), we replaced mCherry with iRFP_670_^41^ (Cry2_olig_-iRFP_670_-NES-NbGFP, Cry2_olig_-iRFP_670_-FUS_N_-NES-NbGFP). Surprisingly, this exchange substantially decreased, both total reporter activation and fold induction. Especially the FUS_N_-containing construct showed around 1.8-fold higher background levels in the reporter assays (Supplementary Figure 2A). By microscopical analysis, we observed the formation of large clusters in darkness, likely leading to this increased background (Supplementary Figure 2B). We concluded, that different fluorescent proteins can influence the clustering behavior of our constructs (similar to previous reports^17^) and thereby the capability of our system to activate the NF-κB pathway. In the case of iRFP_670,_ this is likely due to its tendency to form homodimers^41^. Therefore, we went back to the original, mCherry-tagged constructs for further experiments and tested the impact of the three clustering-increasing constructs.

Interestingly, all three constructs potently increased the signal output of the constructs containing only Cry2 (Cry2-mCh-NES-NbGFP, Cry2-mCh-FUS_N_-NES-NbGFP), both, in terms of fold NF-κB induction (up to 2.8x increase) (Figure 2C) and total NF-κB activity (up to 3.3x increase) (Supplementary Figure 3A). Their activity was now comparable to the Cry2_olig_-mCh-NES-NbGFP condition but did not exceed it (Figure 2C, Supplementary Figure 3). In contrast, for the constructs that already contained Cry2_olig_ (Cry2_olig_-mCh-NES-NbGFP, Cry2_olig_-mCh-FUS_N_-NES-NbGFP), total NF-κB activity increased up to 1.9-fold (Supplementary Figure 3A) but the fold NF-κB induction was not or only slightly improved (maximum 1.4x, Figure 2C). This resulted likely from increased background levels, especially after the addition of the FUS_N_-containing constructs (Supplementary Figure 3B-D). Indeed, we observed the formation of some clusters in darkness, in the FUS_N_-containing conditions that could cause this increased background (Supplementary Figure 4).

**Figure 3:**
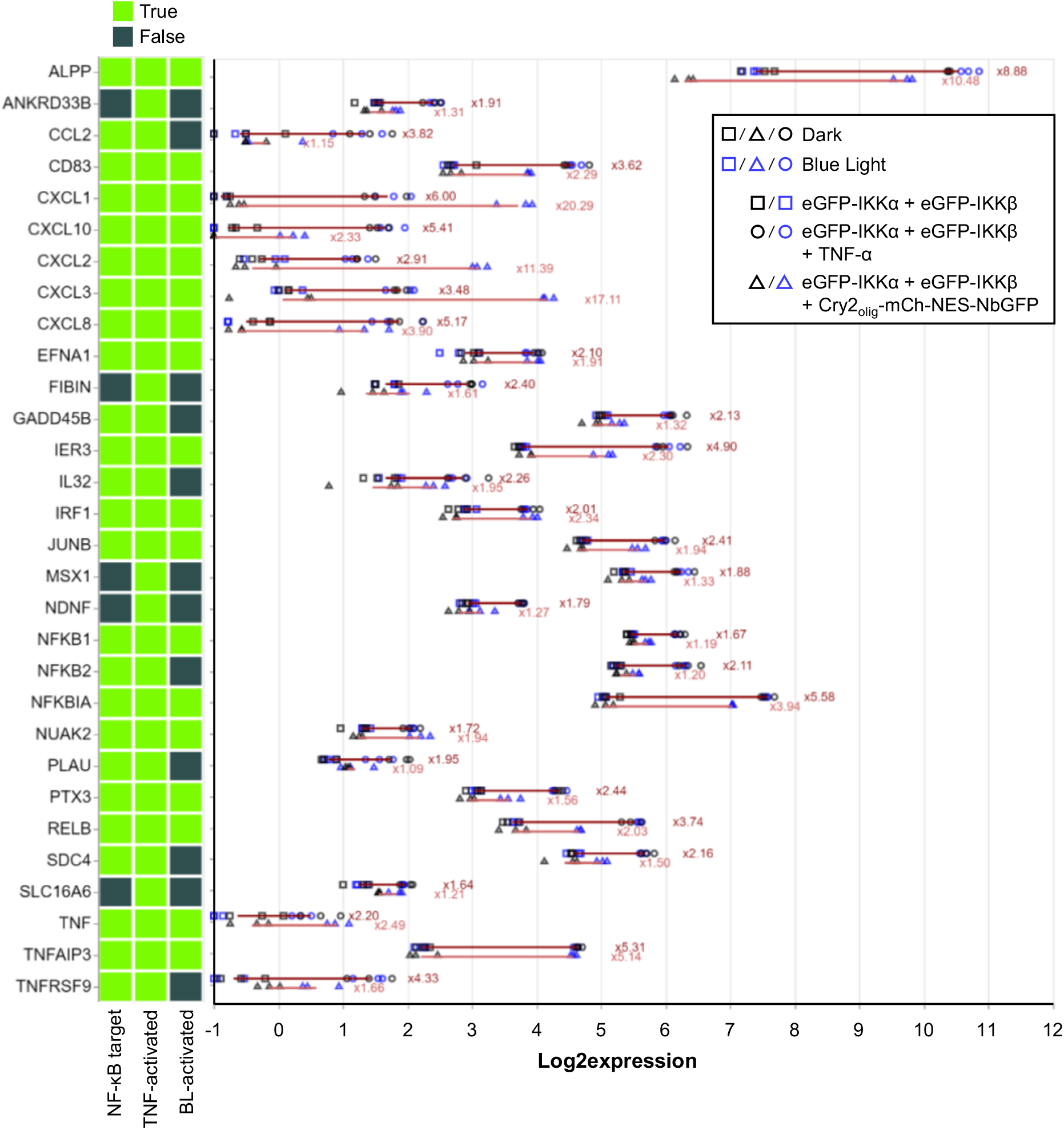
Optogenetic activation of endogenous NF-κB target genes. HEK-293T cells were transfected with the indicated constructs. 24 h after transfection, indicated samples were stimulated with 20 ng/ml TNF-α. Simultaneously, blue-light illumination (5 μmol m^-2^ s^-1^) was started. 3 h later, cells were lysed, and total RNA was extracted and subjected to RNAseq (N = 3 replicates per sample). Genes with a fold activation > 1.61 and a Benjamini-Hochberg-corrected p-value (false discovery rate) < 0.05 between the means of ± TNF-α or D/BL conditions were considered significantly upregulated. Genes marked with false (dark green) in the NF-κB target category are secondary targets. Log2 expression for each gene and replicate is shown. Red lines indicate the fold induction between the means, numbers are the fold inductions.

In summary, the addition of Cry2_olig_-mCh had the highest impact on the systems activity, and we concluded, that clustering of IKK α and β is sufficient to activate NF-κB signaling and that increasing the order of clustering can amplify pathway activation to a certain degree.

### Activation of endogenous NF-κB target genes

After successfully establishing our optogenetic toolbox for NF-κB-specific reporter gene expression, we investigated if our system could also activate endogenous NF-κB target genes. We selected Cry2_olig_-mCh-NES-NbGFP for eGFP-IKKα and β clustering and compared the capability of this configuration to activate NF-kB signaling in comparison with TNF-α stimulation in HEK-293T cells. We extracted total RNA after 3 h of TNF-α stimulation or 3 h of blue-light treatment and analyzed gene expression in the samples via RNA sequencing. We found 30 genes significantly upregulated by TNF-α stimulation and 18 genes significantly upregulated by blue-light illumination. All genes were previously reported to be targets of NF-κB signaling or secondary targets^42–45^, indicating that our blue-light inducible tool is indeed capable of specifically activating NF-κB target genes. Notably, not all TNF-α induced genes were also activated by blue light. However, we observed that these genes were not strongly activated by TNF-α either or secondary targets. The fold induction of many genes was slightly lower for blue light than for TNF-α, therefore blue light mediated activation might not have been strong enough in these cases or, for the example of the secondary targets, not long enough to be activated over an intermediate step. Alternatively, this might be the result of slightly different activation kinetics that could result in maximum expression of different genes to different time points (Figure 3). We furthermore verified these results by analyzing three NF-κB target genes (*NFKBIA, TNFAIP3* and *CXCL1*), via RT-qPCR. Cry2_olig_-mCh-NES-NbGFP induced clustering of eGFP-IKKα and eGFP-IKKβ activated all three target genes, to a comparable or even higher extent than TNF-α stimulation (Supplementary Figure 5).

In conclusion, the here-presented system is capable of potently and specifically activating NF-κB signaling by blue-light induced clustering of IKKα and β, independent from upstream receptor stimulation.

## Discussion

The spatial and temporal organization of the cell is of fundamental importance for all biological processes, including the regulation of signaling pathways. The importance of protein oligomerization, scaffolds, or other mechanisms that generate multivalency has come to special interest as many protein functions rely on the precisely regulated formation of higher-order assemblies and of phase separation^46,47^. Furthermore, signal transduction is not a linear process, but rather a network of tightly regulated signaling patterns that is integrated into dynamic outputs^48^. The gene expression output of the NF-κB pathway differs depending on the inflammatory stimuli: TNF-α treatment leads to the expression of inflammatory genes^49,50^, whereas LPS stimulation induces the expression of genes required for the adaptive immune response^51,52^. Investigation of the differences in signaling patterns that lead to this differential output requires the development of sophisticated, well-controllable, and fine-tunable research tools, that can be provided by optogenetics^53,54^. In comparison to other pathways such as the MAPK pathway, NF-κB signaling has only recently moved into the focus of optogenetic control. DeFelice *et al*. have showcased the potential of optogenetics by engineering Opto-MyD88 and OptoTRAF6, which act downstream of toll-like receptor or interleukin-1 receptor-induced signaling, for light-inducible, Cry2_olig_-mediated clustering^55^. Thereby they could identify IRAK1 as an input dosage sensor for toll-like receptor and interleukin-1 receptor mediated NF-κB signaling^28,56^. Osimiri *et al*. used a combination of the LOVTRAP and an AsLOV-based nuclear import tool to directly shuttle the RelA between the cytoplasm and nucleus, mimicking the oscillating dynamics of the transcription factor and determining the resulting downstream gene activation patterns^57^.

The IKK complex is, in terms of pathway order, positioned in between the two just mentioned approaches and a component downstream of all major NF-κB signaling-inducing receptors. IKKα and β are activated by phosphorylation in their T-loops. For this process, two mechanisms and their relative importance are still discussed^58^: First, phosphorylation by the upstream kinase TAK1^59,60^ and second, auto-transphosphorylation between IKK α and β upon close spatial interaction^61,62^. Recently, it has been shown that both, the IKK complex and TAK1 are recruited into NEMO/polyUb biomolecular condensates, which supports, according to Du *et al*., IKK phosphorylation by TAK1^30^. However, Bagnéris *et al*. recently showed, how the viral oncoprotein vFLIP is sufficient to activate the IKK complex by stabilizing a multimeric configuration that brings the IKKs in close proximity and enables activation by trans-autophosphorylation^63^. Furthermore, it has been shown that enforced homooligomerization of IKKα or β via three repeats of the chemical dimerizer FPK is capable of activating NF-κB, as well^64^. Cry2-induced clustering could facilitate a similar mechanism but full activation might not be achieved due to the lack of TAK1-induced phosphorylation or other unknown mechanisms.

In contrast to RelA overexpression, the IKKs are mainly inactive in the here-used over-expression state, enabling the OFF state of the system. Our here-developed optogenetic toolbox enables testing of the optimal oligomerization state of the IKK signaling module. We linked the different oligomerization states to differently high signal outputs and found that there seems to be a capacity limit for the IKKs, above which more clustering is not beneficial anymore. Furthermore, phase separation in comparison to efficient clustering, increased the activation strength only marginally and the addition of Cry2_olig_-mCh-FUS_N_-NES led to an increased background in the dark state. Additionally, our RNAseq analysis showed that the system is capable of specifically activating a set of endogenous NF-κB target genes.

In conclusion, the here-developed optogenetic toolbox enables the systematic testing for the optimal homo-or heteromeric clustering state of a target protein. It should be considered that the optimum may vary significantly depending on the target’s intrinsic properties, its function in the pathway, or the desired goal of the user. The constructs could be used in the future to test different input lengths, strengths, or frequencies simply by switching blue light ON or OFF. Therefore, the toolbox represents a sophisticated approach to decipher the signaling dynamics of NF-κB or other pathways, simply by fusing the clustering-regulated components to eGFP.

## Materials and Methods

### Cloning

All plasmids generated in this study were cloned with Gibson Assembly^65^ or Aqua Cloning^66^. Detailed plasmid descriptions are listed in Table S1. Linkers were inserted with oligonucleotides and PCR. For constructs that contain multiple repeats, such as response elements in promoters, *E. coli* were grown at 30 °C. All plasmids were verified with Sanger sequencing.

### Cell culture and transfection

HEK-293T cells (DSMZ, catalog no. ACC 635) were cultivated at 37 °C and 5% CO_2_ in Dulbecco’s modified Eagle’s medium (DMEM, PAN Biotech, catalog no. P04-03550) supplemented with 10% (v/v) fetal calf serum (FCS, PAN Biotech, catalog no. P30-3602) and 1% (v/v) penicillin-streptomycin solution (PAN Biotech, catalog no. P06-07100). Cells were passaged when reaching a confluency of ≈ 80%, every 2-3 days. All experiments were conducted in multi-well plates as described below. Cells were transfected 24 h after seeding with polyethyleneimine (PEI, 1 mg/ml in H_2_O, pH 7, Polyscience, catalog no. 23966-1). To do so, the total volume of Opti-MEM (Thermo Fisher Scientific, catalog no. 22600-134) was split into two mixes: DNA-mix and PEI-mix (exact amounts are described below for each experiment scale), which were prepared separately. Afterwards, the two solutions were combined, vortexed immediately for 15 s, and then incubated for 15 min at room temperature. DNA/PEI mixes were then applied to the cells dropwise. For optogenetic experiments, cells were then kept in darkness. Plasmid amounts and combinations for each experiment are described in Table S2.

### Reporter Assays

Firefly luciferase assays were performed in black, clear-bottom 96-well plates (μCLEAR®, Greiner Bio-One, catalog no. 655090). 15,000 cells/well in 100 μl DMEM were seeded 24 h prior to transfection. Transfection mixes contained 125 ng DNA and 0.41 μl PEI in 20 μl Opti-MEM per well. Illumination (5 μmol m^-2^ s^-1^) of optogenetic experiments was started 8 h after transfection using the optoPlate-96^67^ equipped with 470 nm LEDs (Würth Elektronik, MPN: 150141RB73100). The optoPlate-96 was programmed using the optoConfig-96^68^. Stimulation of the controls was performed 8 h after transfection with 20 ng/ml TNF-α (Merck, cat. no.: H8916). Cells were lysed 24 h later by addition of 100 μl lysis buffer (25 mM Tris/HCl, pH 7.8, 1% Triton X-100, 15 mM MgSO_4_, 4 mM ethylene glycol tetraacetic acid (EGTA), 1 mM DTT) per well and 5 min incubation at room temperature (RT). Cells were resuspended by pipetting up and down carefully and 40 μl lysate of each well was transferred into a white flat-bottom 96-well plates (Corning Incorporated, catalog no. CORN3912). Afterwards, the plate was centrifuged for 30 s at 1200 rpm. To measure firefly luciferase activity 20 μl firefly substrate (20 mM Tricine, 2.67 mM MgSO_4_, 0.1 mM EDTA, 33.3 mM DTT, 0.52 mM ATP, 0.27 mM Acetyl-CoA, 5 mM NaOH, 0.264 mM MgCO_3_, 0.47 mM luciferin) per well was added. The measurement was started immediately using either a Synergy 4 multimode microplate reader (BioTek Instruments Inc.) or a SpectraMax iD5 microplate reader (Molecular Devices GmbH). Program: 10 s shaking, 1000 ms integration time, endpoint measurement. For the calculation of fold induction or fold signal amplification, only samples that were grown and measured on the same plate were compared.

### Microscopy

For microscopy experiments cells were grown on collagen-coated glass coverslips (Carl Roth, catalog no.: YX03.2), in 24-well format (Corning, catalog no. CORN3524). For coating, 500 μl rat tail collagen I (50 μg/ml diluted in 25 mM acetic acid, Thermo Fisher Scientific, catalog no. A1048301) was added to each well for one hour at RT. Then, wells were washed three times with 500 μl PBS, and 75,000 cells per well were seeded. Transfection mixes for 24-well format contained: 750 ng DNA and 2.4 μl PEI in 100 μl OptiMEM per well. Illumination (5 μmol m^-2^ s^-1^) was started 8 h after transfection for 24 h in ventilated boxes containing microcontroller-regulated illumination panels with 460 nm LEDs (LED Engin, MPN: LZ1-10B202-0000). Controls were kept in identical boxes but in darkness. For fixation, the medium was removed and 200 μl methanol-free paraformaldehyde (PFA, Science Services, catalog no. E15714-S; 4% diluted in PBS v/v) was added under dim, green safe light. The samples were incubated for 15 min at RT and in darkness, then PFA was removed and cells were washed twice with 500 μl PBS. For the staining of nuclei, 500 μl 0.2 μg/ml 4’,6’-diamidino-2-phenylindole (DAPI, Merck, catalog no. D9542) in PBS was added for 15 min at RT. Cells were washed twice afterwards with 500 μl PBS and mounted on microscopy slides onto 8.5 μl Mowiol mounting medium (2.4 g of Mowiol, 6 g of glycerol, 6 ml of H_2_O and 12 ml of Tris/HCl (pH 8.5)). Samples were dried and then additionally fixed with transparent nail polish. Images were acquired with a Zeiss LSM 880 laser scanning confocal microscope equipped with a 63x Plan-Apochromat oil objective (NA 1.4) in z-stacks with 1 μm distance. DAPI was imaged with the 405 nm laser, eGFP with the 488 nm laser, mCherry with the 561 nm laser, and iRFP670 with the 633 nm laser. All images are displayed as cutouts of the maximum intensity projections with equally adjusted intensities.

### RNAseq and reverse transcription quantitative PCR (RT-qPCR)

For RNAseq and RT-qPCR experiments, 90,000 HEK-293T cells were seeded into 24-well plates, transfected as described above for 24-well format, and kept in darkness for 24 h. Afterwards, cells were illuminated (460 nm, 5 μmol m^−2^ s^−1^) in the boxes described above for 3 h, controls were kept in darkness. Simultaneously, TNF-α (20 ng/ml) was applied to the control group. Afterwards, cells were harvested and RNA was isolated using the RNeasy plus micro kit (Qiagen, cat. no.: 74034), according to the manufacturer’s protocol. RNA integrity was verified with an agarose gel and 260/280 and 230/280 ratios were measured with a nanodrop 1000 (Thermo Fisher Scientific). RNAseq was performed by BGI Tech Solutions (Hong Kong) Co. after where RNA integrity was again verified with a Bioanalyzer (Agilent 2100 Bioanalyzer).

For RT-qPCR cDNA was generated using the High Capacity Kit (Applied Biosystems™, cat. no. 4368814) according to the manufacturer’s protocol using 1 μg RNA, cDNA was diluted 1:3.

Primer sequences for the three NF-κB target genes were chosen from literature: (*NFK-BIA*, oAF469: 5’-ATGTCAATGCTCAGGAGCCC-3’ and oAF470: 5’-GACATCAGCCCCACACTTCA-3’^69^; *TNFAIP3*, oAF467: 5’-CGTCCAGGTTCCAGAACACCATTC-3’ and oAF468: 5’-TGCGCTGGCTCGATCTCAGTTG-3’^70^; *CXCL1*, oAF471: 5’-AAC-CGAAGTCATAGCCACAC-3’ and oAF472: 5’-GTTGGATTTGTCACTGTTCAGC-3’^71^).

*GUS* was selected as housekeeping gene for normalization (oMH703/oAF481, 5’-CGTCCCACCTAGAATCTGCT-3’ and oMH702/oAF482, 5’-

TTGCTCACAAAGGTCACAGG-3’). qPCRs were conducted in technical triplicates, in 10 μl approaches using the PowerTrack™ SYBR Green Mastermix (Applied Biosys-tems™, cat. no. A46110) according to the manufacturer’s protocol (fast cycling mode with default dissociation step). A CFX384 Touch Real-Time PCR Detection System (BioRad) was used for the measurement. Data was analyzed with the ΔΔCt method and is displayed as fold changes to the negative controls.

### RNAseq analysis

The externally provided raw but already quality-controlled and cleaned data in FASTQ format was analyzed using a method based on gene-specific gapped k-mer counting. A unique gapped k-mer is a discontiguous DNA sequence of length k (here, k=23 in a window of length w=33 using the specific mask ######_##_### # ###_##_######, where # and _ denote the 23 considered and 10 ignored positions in a window, respectively) that is specific for a gene in the t2t reference genome^72^, i.e., occurs nowhere else in the genome than in that gene. The set of such unique gapped k-mers was first identified using the hackgap k-mer counter^73^. The number of occurrences of each such k-mer in each FASTQ sample file was then counted, again using hackgap. Per gene and per sample, k-mer counts were robustly averaged using trimmed means to yield a measure of gene expression. The resulting count matrix was normalized by a scaling factor to account for slightly varying sequencing depth between samples using quantile regression. In detail, all quantiles between median (0.5-quantile) and 0.98-quantile of the sorted expression values of each sample were averaged over all samples to yield an average quantile curve. Then, the optimal scaling factor (offset in logarithmic space) was computed for each sample to regress the samples’ quantiles onto the average curve. The resulting normalized expression matrix was subjected to statistical testing and fold change estimation fold changes were estimated as ratios between geometric means of the three replicates in each condition. The compared conditions were -/+ TNF-α on the one hand and darkness vs. blue light (D/BL) on the other hand.

Statistical tests were performed with the scipy.stats^74^ package (v1.12) of Python (v3.11.7), using a two-sample t-test with unequal variance, combined with Benjamini-Hochberg p-value multiple testing correction for false discovery rate (FDR) control (functions ttest_ind, false_discovery_control). The whole analysis workflow was implemented with Snakemake^75,76^.

Genes with a fold change > 1.61 and a false discovery rate < 0.05 between the means of -/+ TNF-α or D/BL conditions were considered significantly upregulated. NF-κB targets and secondary targets were determined according to the list of NF-κB target genes provided by the Gilmore Laboratory (Boston University)^77^, the GeneCards database^78^, and the Enrichr gene list enrichment analysis tool^79,80^.

### Other software

Analyses were performed with Microsoft Excel 2016 and graphs were generated with GraphPad Prism 9.2.0. Images were analyzed with ImageJ 2.3.0/1.53q including the Biovoxxel Toolbox v2.6.0^81^. FACs data was analyzed using FlowJo-v10™. All schemes were created with BioRender.com.

## Acknowledgments

We are very grateful to M. Klenzendorf and D. Gaspar for their excellent technical assistance. We would like to thank the Faculty of Biology technical workshop for the construction of the illumination devices. We acknowledge the excellent scientific and technical assistance of the Signalling Factory Core Facility staff of the Albert-Ludwigs-University Freiburg, especially P. Salavei and N. Gensch. We acknowledge the staff of the Life Imaging Center (LIC) in the Hilde Mangold Haus (HMH) of the Albert-Ludwigs-University Freiburg for their help with microscopy resources and their excellent support in microscopy setup and image acquisition.

## Author contributions

A.A.M.F. and M.M.G. performed experimental work. A.A.M.F., S.R. and W.W. analyzed the results and wrote the manuscript. S.R. implemented the RNA-seq analysis method. M.F. and B.G. contributed to the design of the study and data interpretation.

W.W. supervised the work.

## Data and materials availability

All data needed to evaluate the conclusions in the paper is present in the paper and/or Supplementary Materials. Raw data (images, luciferase activity measurements) are available from the authors upon request. Plasmids and detailed plasmid maps are available from the authors upon request.

## References

1. Garratt, R. C., Valadares, N. F. & Bachega, J. F. R. Oligomeric Proteins. Encyclopedia of Biophysics 1781–1818 (2013).

2. Ali, M. H. & Imperiali, B. Protein oligomerization: How and why. Bioorg Med Chem 13, 5013–5020 (2005).

3. Hashimoto, K., Nishi, H., Bryant, S. & Panchenko, A. R. Caught in self-interaction: Evolutionary and functional mechanisms of protein homooligomerization. Phys Biol 8, 035007 (2011).

4. Kholodenko, B. N., Hancock, J. F. & Kolch, W. Signalling ballet in space and time. Nat Rev Mol Cell Biol 11, 414–426 (2010).

5. Kennedy, M. J. et al. Rapid blue light induction of protein interaction in living cells. Nat Methods 7, 973–975 (2010).

6. Kawano, F., Suzuki, H., Furuya, A. & Sato, M. Engineered pairs of distinct photoswitches for optogenetic control of cellular proteins. Nat Commun 6, 6256 (2015).

7. Wang, X., Chen, X. & Yang, Y. Spatiotemporal control of gene expression by a light-switchable transgene system. Nat Methods 9, 266–271 (2012).

8. Guntas, G. et al. Engineering an improved light-induced dimer (iLID) for controlling the localization and activity of signaling proteins. PNAS 112, 114–117 (2015).

9. Levskaya, A., Weiner, O. D., Lim, W. A. & Voigt, C. A. Spatiotemporal control of cell signalling using a light-switchable protein interaction. Nature 461, 997–1001 (2009).

10. Bugaj, L. J., Choksi, A. T., Mesuda, C. K., Kane, R. S. & Schaffer, D. V. Optogenetic protein clustering and signaling activation in mammalian cells. Nat Methods 10, 249–254 (2013).

11. Kainrath, S., Stadler, M., Reichhart, E., Distel, M. & Janovjak, H. Green-Light-Induced Inactivation of Receptor Signaling Using Cobalamin-Binding Domains. Angewandte Chemie International Edition 56, 4608–4611 (2017).

12. Masuda, S., Nakatani, Y., Ren, S. & Tanaka, M. Blue light-mediated manipulation of transcription factor activity in vivo. ACS Chem Biol 8, 2649–2653 (2013).

13. Kolar, K., Knobloch, C., Stork, H., Žnidarič, M. & Weber, W. OptoBase: A Web Platform for Molecular Optogenetics. ACS Synth Biol 7, 1825–1828 (2018).

14. Shin, Y. et al. Spatiotemporal Control of Intracellular Phase Transitions Using Light-Activated optoDroplets. Cell 168, 159-171.e14 (2017).

15. Bracha, D. et al. Mapping Local and Global Liquid Phase Behavior in Living Cells Using Photo-Oligomerizable Seeds. Cell 175, 1467-1480.e13 (2018).

16. Taslimi, A. et al. An optimized optogenetic clustering tool for probing protein interaction and function. Nat Commun 5, 4925 (2014).

17. Park, H. et al. Optogenetic protein clustering through fluorescent protein tagging and extension of CRY2. Nat Commun 8, 1–7 (2017).

18. Kramer, M. M., Lataster, L., Weber, W. & Radziwill, G. Optogenetic approaches for the spatiotemporal control of signal transduction pathways. Int J Mol Sci 22, (2021).

19. Mühlhäuser, W. W. D., Fischer, A., Weber, W. & Radziwill, G. Optogenetics - Bringing light into the darkness of mammalian signal transduction. Biochimica et Biophysica Acta - Molecular Cell Research vol. 1864 280–292 Preprint at 10.1016/j.bbamcr.2016.11.009 (2017).

20. Wend, S. et al. Optogenetic control of protein kinase activity in mammalian cells. ACS Synth Biol 3, 280–285 (2014).

21. Chatelle, C. V. et al. Optogenetically controlled RAF to characterize BRAF and CRAF protein kinase inhibitors. Sci Rep 6, 23713 (2016).

22. Aoki, H. & Hiratsuka, T. Propagating Wave of ERK Activation Orients Collective Cell Migration Article Propagating Wave of ERK Activation Orients Collective Cell Migration. Dev Cell 43, 305-317.e5 (2017).

23. Ender, P. et al. Spatiotemporal control of ERK pulse frequency coordinates fate decisions during mammary acinar morphogenesis. Dev Cell 57, 2153–2167 (2022).

24. Repina, N. A. et al. Optogenetic control of Wnt signaling models cell-intrinsic embryogenic patterning using 2D human pluripotent stem cell culture. Hum Dev 150, dev201386 (2023).

25. Leopold, A. V. & Verkhusha, V. V. Light control of RTK activity: From technology development to translational research. Chem Sci 11, 10019–10034 (2020).

26. Bugaj, L. J. et al. Regulation of endogenous transmembrane receptors through optogenetic Cry2 clustering. Nat Commun 6, (2015).

27. Kim, N. et al. Resource Spatiotemporal Control of Fibroblast Growth Factor Receptor Signals by Blue Light. Chem Biol 21, 903–912 (2014).

28. DeFelice, M. M. et al. NF-κB signaling dynamics is controlled by a dose-sensing autoregulatory loop. Sci Signal 12, eaau3568 (2019).

29. Wu, H. Higher-order assemblies in a new paradigm of signal transduction. Cell 153, 287–292 (2013).

30. Du, M., Ea, C. K., Fang, Y. & Chen, Z. J. Liquid phase separation of NEMO induced by polyubiquitin chains activates NF-κB. Mol Cell 82, 2415-2426.e5 (2022).

31. Goel, S. et al. Linear ubiquitination induces NEMO phase separation to activate NF-κB signaling. Life Sci Alliance 6, 1–18 (2023).

32. Hinz, M. & Scheidereit, C. The IκB kinase complex in NF-κB regulation and beyond. EMBO Rep 15, 46–61 (2014).

33. Chen, Z. J. Ubiquitination in signaling to and activation of IKK. Immunol Rev 246, 95–106 (2012).

34. Israël, A. The IKK complex, a central regulator of NF-kappaB activation. Cold Spring Harb Perspect Biol 2, 1–15 (2010).

35. Wajant, H. & Scheurich, P. TNFR1-induced activation of the classical NF-κB pathway. FEBS Journal 278, 862–876 (2011).

36. Kirchhofer, A. et al. Modulation of protein properties in living cells using nanobodies. Nat Struct Mol Biol 17, 133–139 (2009).

37. Lee, S. et al. Reversible protein inactivation by optogenetic trapping in cells. Nat Methods 11, 633–636 (2014).

38. Ochoa-Fernandez, R. et al. Optogenetic control of gene expression in plants in the presence of ambient white light. Nat Methods 17, 717–725 (2020).

39. Mahlandt, E. K., Toereppel, M., haydary, T. & Goedhart, J. iLID-antiGFP-nanobody is a flexible targeting strategy for recruitment to GFP-tagged proteins. bioRxiv (2022).

40. Schneider, P., Willen, L. & Smulski, C. R. Tools and techniques to study ligand-receptor interactions and receptor activation by TNF superfamily members. Methods Enzymol 545, 103–125 (2014).

41. Shcherbakova, D. M. & Verkhusha, V. V. Near-infrared fluorescent proteins for multicolor in vivo imaging. Nat Methods 10, 751–754 (2013).

42. Boston University. NF-κB Gene Resources: Target Genes. https://www.bu.edu/nf-kb/gene-resources/target-genes/ (2023).

43. G, S. et al. The GeneCards Suite: From Gene Data Mining to Disease Genome Sequence Analyses. Curr Protoc Bioinformatics 54, 1.30.1-1.30.33 (2016).

44. Chen, E. Y. et al. Enrichr: interactive and collaborative HTML5 gene list enrichment analysis tool. BMC Bioinformatics 14, 128 (2013).

45. Kuleshov, M. V. et al. Enrichr: a comprehensive gene set enrichment analysis web server 2016 update. Nucleic Acids Res 44, W90–W97 (2016).

46. Good, M. C., Zalatan, J. G. & Lim, W. A. Scaffold Proteins : Hubs for Controlling the Flow of Cellular Information Domains. Science (1979) 332, 680–687 (2011).

47. Su, Q., Mehta, S. & Zhang, J. Liquid-liquid phase separation : Orchestrating cell signaling through time and space. Mol Cell 81, 4137–4146 (2021).

48. Purvis, J. E. & Lahav, G. Encoding and decoding cellular information through signaling dynamics. Cell 152, 945–956 (2013).

49. Hoffmann, A., Levchenko, A., Scott, M. L. & Baltimore, D. The IκB-NF-κB signaling module: Temporal control and selective gene activation. Science (1979) 298, 1241–1245 (2002).

50. Nelson, D. E. et al. Oscillations in NF-κB signaling control the dynamics of gene expression. Science (1979) 306, 704–708 (2004).

51. Covert, M. W., Leung, T. H., Gaston, J. E. & Baltimore, D. Achieving stability of lipopolysaccharide-induced NF-κB activation. Science (1979) 309, 1854–1857 (2005).

52. Werner, S. L., Barken, D. & Hoffmann, A. Stimulus specificity of gene expression programs determined by temporal control of IKK activity. Science (1979) 309, 1857–1861 (2005).

53. Fischer, A. A. M., Kramer, M. M., Radziwill, G. & Weber, W. Shedding light on current trends in molecular optogenetics. Curr Opin Chem Biol 70, 102196 (2022).

54. Farahani, P. E., Reed, E. H., Underhill, E. J., Aoki, K. & Toettcher, J. E. Signaling, Deconstructed: Using Optogenetics to Dissect and Direct Information Flow in Biological Systems. Annu Rev Biomed Eng 23, 61–87 (2021).

55. DeFelice, M. M. et al. NF-κB signaling dynamics is controlled by a dose-sensing auto-regulatory loop. Sci Signal 12, eaau3568 (2019).

56. Wu, M. et al. Negative regulators of STAT3 signaling pathway in cancers. Cancer Manag Res 11, 4957–4969 (2019).

57. Osimiri, L. C. et al. Optogenetic control of RelA reveals effect of transcription factor dynamics on downstream gene expression. bioRxiv (2022).

58. Hayden, M. S. & Ghosh, S. NF-kB , the first quarter-century : remarkable progress and outstanding questions. Genes Dev 26, 203–234 (2012).

59. Ninomiya-Tsuji, J. et al. The kinase TAK1 can activate the NIK-IκB as well as the MAP kinase cascade in the IL-1 signalling pathway. Nature 398, 252–256 (1999).

60. Chen Wang et al. TAK1 is a ubiquitin-dependent kinase of MKK and IKK. Nature 412, 346–351 (2001).

61. Polley, S. et al. A Structural Basis for IκB Kinase 2 Activation Via Oligomerization-Dependent Trans Auto-Phosphorylation. PLoS Biol 11, e1001581 (2013).

62. Delhase, M., Makio, H., Chen, Y. & Karin, M. Positive and negative regulation of IκB kinase activity through IKKβ subunit phosphorylation. Science (1979) 284, 309–313 (1999).

63. Bagnéris, C. et al. Mechanistic insights into the activation of the IKK kinase complex by the Kaposi’s sarcoma herpes virus oncoprotein vFLIP. Journal of Biological Chemistry 298, 102012 (2022).

64. Inohara, N. et al. An Induced Proximity Model for NF-kB Activation in the Nod1 / RICK and RIP Signaling Pathways. J Biol Chem 275, 27823–27831 (2000).

65. Gibson, D. G. et al. Enzymatic assembly of DNA molecules up to several hundred kilobases. Nat Methods 6, 343–345 (2009).

66. Beyer, H. M. et al. AQUA Cloning: A Versatile and Simple Enzyme-Free Cloning Approach. PLoS One 10, e0137652 (2015).

67. Bugaj, L. J. & Lim, W. A. High-throughput multicolor optogenetics in microwell plates. Nat Protoc 14, 2205–2228 (2019).

68. Thomas, O. S., Hörner, M. & Weber, W. A graphical user interface to design high-throughput optogenetic experiments with the optoPlate-96. Nat Protoc 15, 2785–2787 (2020).

69. Callegari, A. et al. Single-molecule dynamics and genome-wide transcriptomics reveal that NF-kB (p65)-DNA binding times can be decoupled from transcriptional activation. PLoS Genet 15, 1–23 (2019).

70. Jiang, X. et al. Expression of tumor necrosis factor alpha-induced protein 3 mRNA in peripheral blood mononuclear cells negatively correlates with disease severity in psori-asis vulgaris. Clinical and Vaccine Immunology 19, 1938–1942 (2012).

71. Wang, D. et al. CXCL1 induced by prostaglandin E2 promotes angiogenesis in colo-rectal cancer. Journal of Experimental Medicine 203, 941–951 (2006).

72. T2T Consortium. Genome assembly T2T-CHM13v2.0. Genome assembly T2T-CHM13v2.0 (2022).

73. Jens Zentgraf, S. R. hackgap. https://gitlab.com/rahmannlab/hackgap (2022).

74. Virtanen, P. et al. SciPy 1.0: fundamental algorithms for scientific computing in Python. Nat Methods 17, 261–272 (2020).

75. Mölder, F. et al. Sustainable data analysis with Snakemake. F1000Res 10, 33 (2021).

76. Köster, J. & Rahmann, S. Snakemake-a scalable bioinformatics workflow engine. Bioinformatics 28, 2520–2522 (2012).

77. Boston University. NF-κB Gene Resources: Target Genes. https://www.bu.edu/nf-kb/gene-resources/target-genes/ (2023).

78. Stelzer, G. et al. The GeneCards Suite: From Gene Data Mining to Disease Genome Sequence Analyses. Curr Protoc Bioinformatics 54, (2016).

79. Chen, E. Y. et al. Enrichr: interactive and collaborative HTML5 gene list enrichment analysis tool. BMC Bioinformatics 14, 128 (2013).

80. Kuleshov, M. V. et al. Enrichr: a comprehensive gene set enrichment analysis web server 2016 update. Nucleic Acids Res 44, W90–W97 (2016).

81. Brocher, J. BioVoxxel Toolbox v2.6.0. https://zenodo.org/record/8214743 https://zenodo.org/record/8214743 (2023).

