## Supplementary Information for "Activation of NF-κB signaling by optogenetic clustering of IKKα and β"

### Table of contents:


| | | |
|---|---|---|
| Supplementary Figure 1: | Activation of NF-κB signaling by TNF-α stimulation | 3 |
| Supplementary Figure 2: | Impact of the fluorescent protein on the clustering behavior | 4 |
| Supplementary Figure 3: | Amplification of pathway activation with an additional clustering construct | 6 |
| Supplementary Figure 4: | Microscopical characterization of the NbGFP clustering constructs with additional $Cry2_{olig}$-mCh-$FUS_N$-NES | 7 |
| Table S1: | Plasmids used in this study | 8 |
| Table S2: | Transfection conditions of each experiment | 9 |

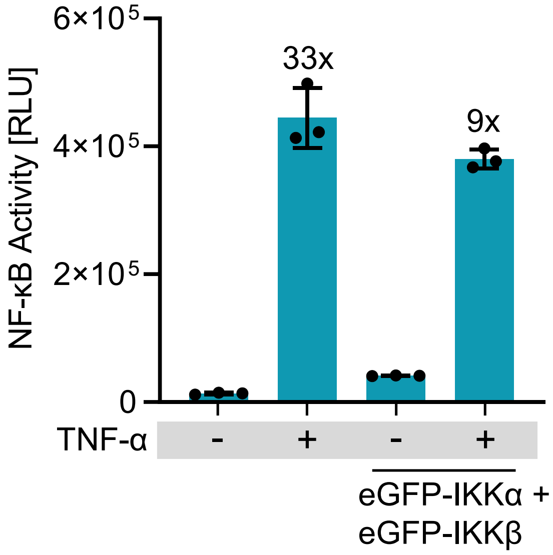


**Supplementary Figure 1: Activation of NF-κB signaling by TNF-α stimulation** HEK-293T cells were transfected with an NF-κB-responsive firefly luciferase reporter and either with an empty vector or eGFP-IKKα and eGFP-IKKβ. 8 h after transfection, indicated samples were stimulated with 20 ng/ml TNF-α. 24 h later, firefly luciferase activity was measured. Mean ± SD and single values are shown (N = 3), and fold NF-κB activation between ± TNF-α is shown above the bar.

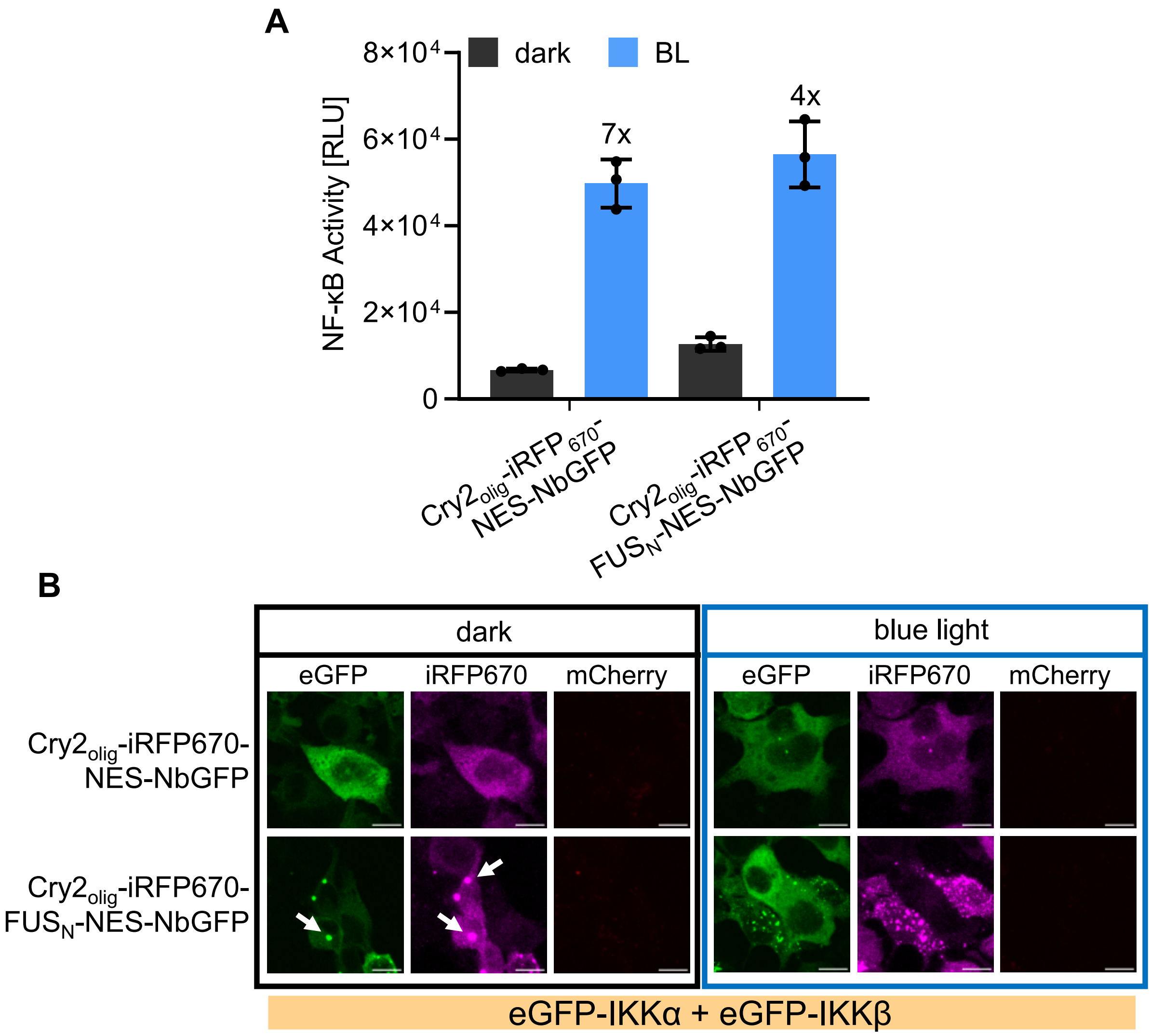


**Supplementary Figure 2: Impact of the fluorescent protein on the clustering behavior**

(**A**) HEK-293T cells were transfected with the indicated constructs and an NF-κB-responsive firefly luciferase reporter. Blue-light illumination (5 µmol $m^{-2}$ $s^{-1}$) was started 8 h after transfection and firefly luciferase activity was determined 24 h later. Mean ± SD and fold NF-κB activation is shown (N = 3). **(B)** Colocalization of the $Cry2_{olig}$-$iRFP_{670}$-NES-NbGFP constructs with $Cry2_{olig}$-mCh-$FUS_N$-NES. HEK-293T cells were treated as described in A but fixed for microscopy analysis. Representative images are shown; scale bar = 10 µm.

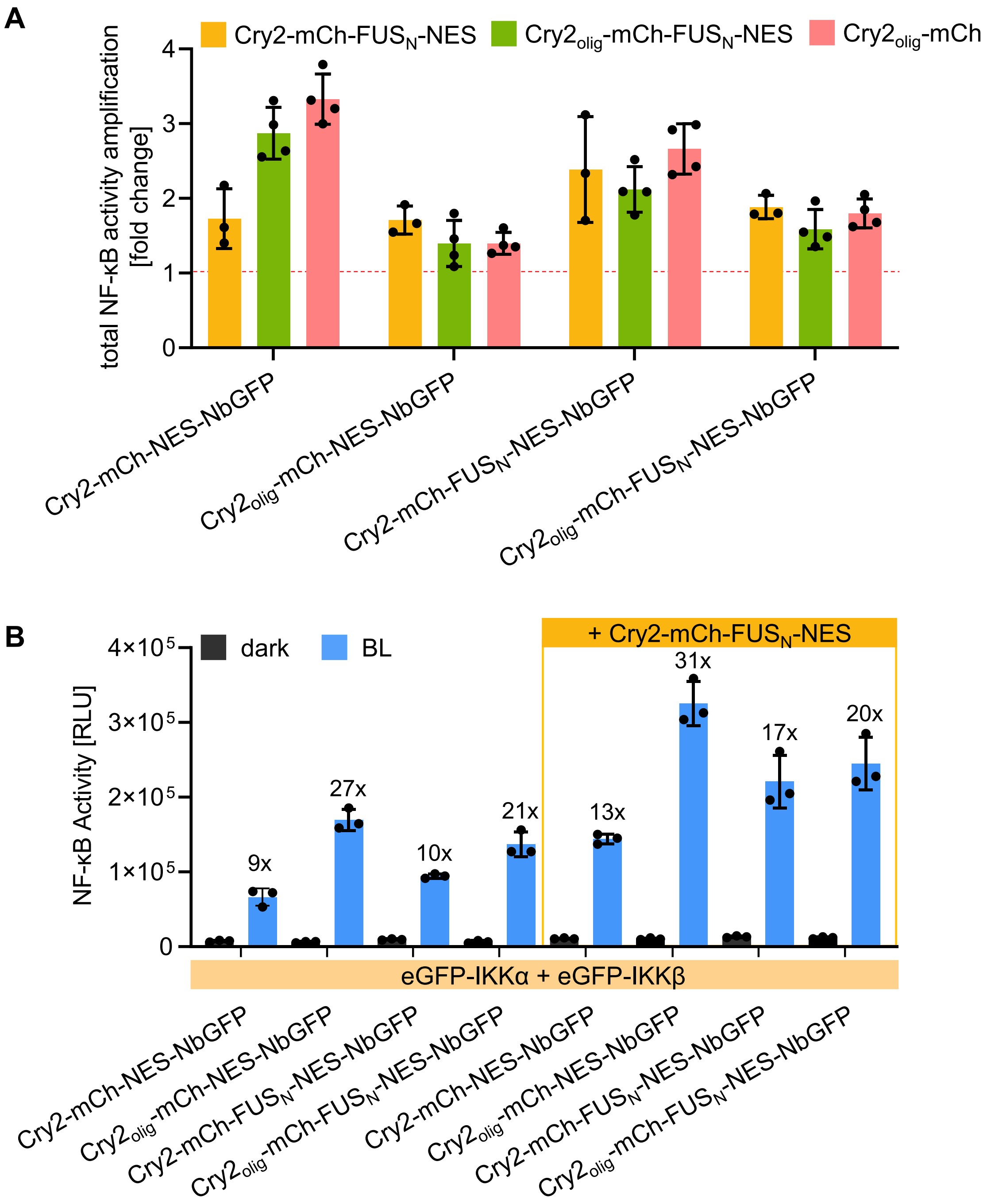
A
Cry2-mCh-FUS$_N$-NES
Cry2$_{olig}$-mCh-FUS$_N$-NES
Cry2$_{olig}$-mCh
total NF-κB activity amplification [fold change]
Cry2-mCh-NES-NbGFP
Cry2$_{olig}$-mCh-NES-NbGFP
Cry2-mCh-FUS$_N$-NES-NbGFP
Cry2$_{olig}$-mCh-FUS$_N$-NES-NbGFP
B
dark
BL
+ Cry2-mCh-FUS$_N$-NES
NF-κB Activity [RLU]
9x
27x
10x
21x
13x
31x
17x
20x
eGFP-IKKα + eGFP-IKKβ

C

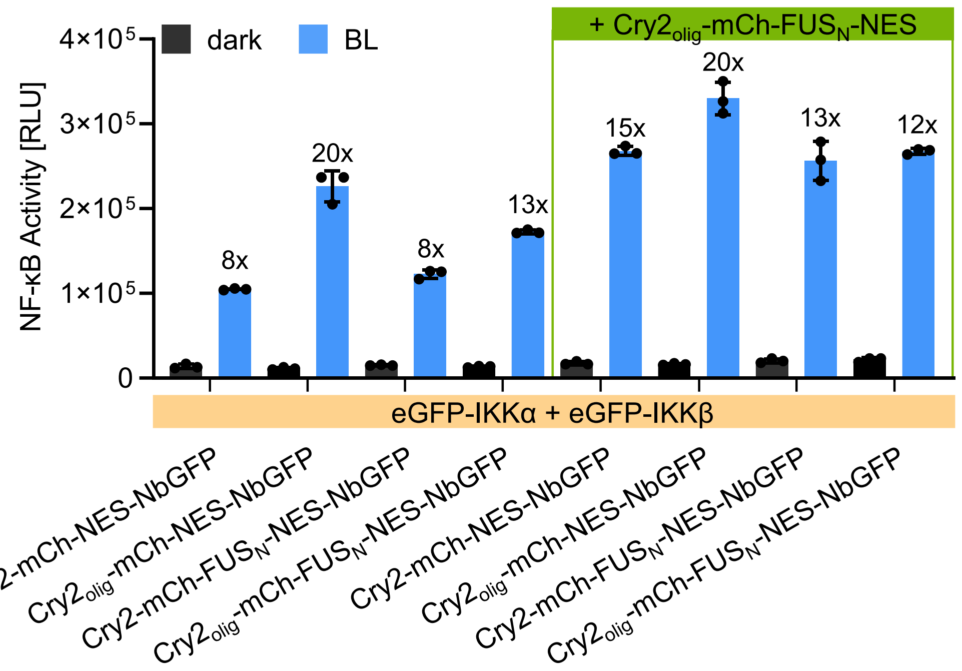


D

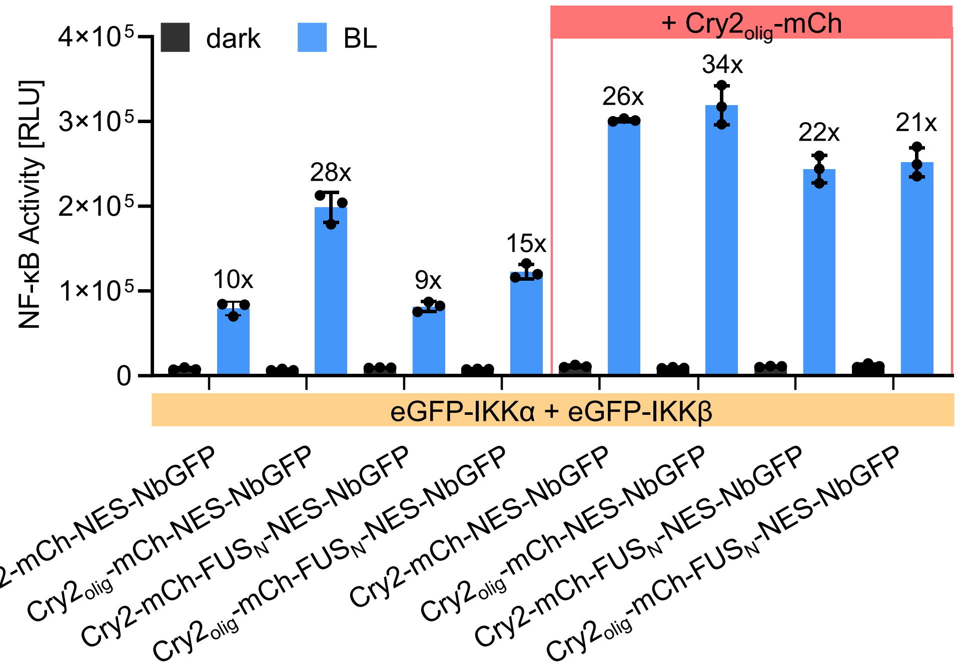


**Supplementary Figure 3: Amplification of pathway activation with an additional clustering construct**

HEK-293T cells were transfected with the indicated constructs and an NF-κB-responsive firefly luciferase reporter. Blue-light illumination (5 µmol $m^{-2}$ $s^{-1}$) was started 8 h after transfection and firefly luciferase activity was measured 24 h later. (**A**) Fold amplification of total NF-κB activity that results from the addition of the indicated additional clustering construct (Cry2-mCh-$FUS_N$-NES, $Cry2_{olig}$-mCh-$FUS_N$-NES, or $Cry2_{olig}$-mCh). Mean ± SD of N = 3-4 independent experiments. (**B-D**) Raw values of blue-light induced NF-κB activity. Mean ± SD and fold NF-κB activation is shown, N = 3 replicates. **Please note:** The first 4 groups in C is the same data as shown in Figure 2B. It is additionally shown here for direct comparison with the + $Cry2_{olig}$-mCh-$FUS_N$-NES condition.

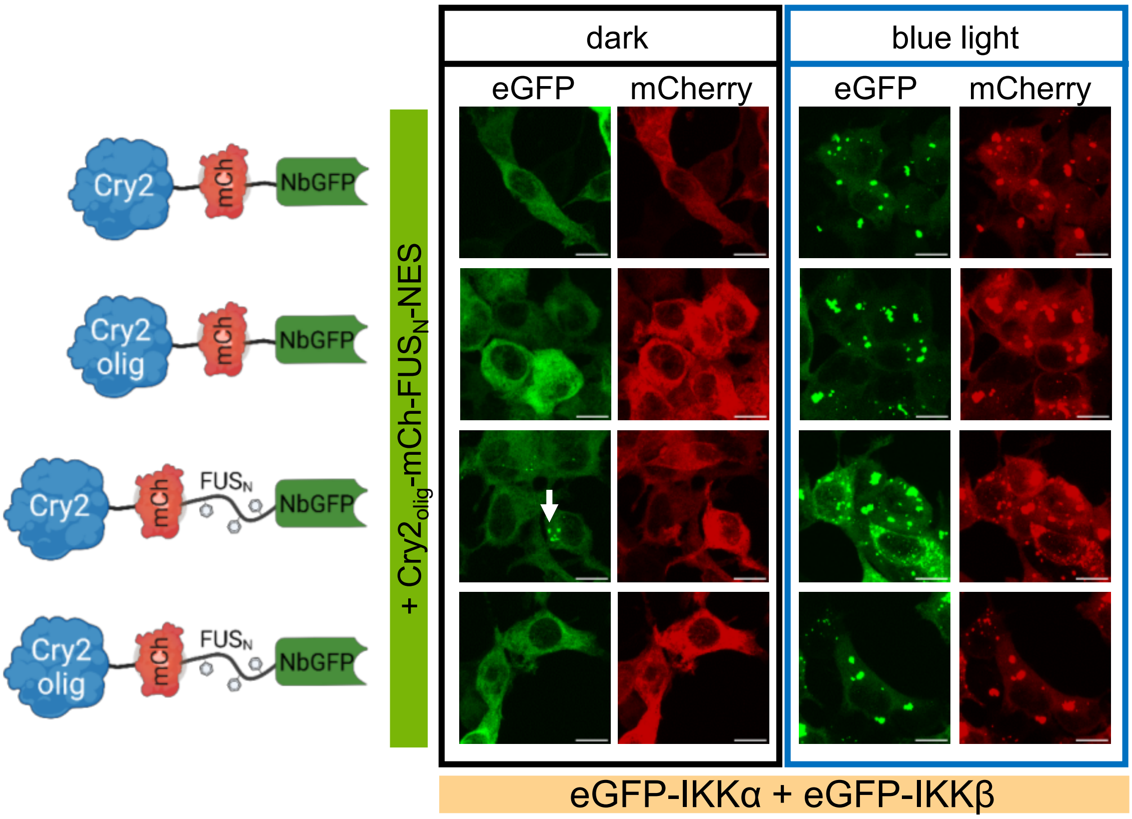


**Supplementary Figure 4: Microscopical characterization of the NbGFP clustering constructs with additional $Cry2_{olig}$-mCh-$FUS_N$-NES**

HEK-293T cells were transfected with the indicated constructs, an NF-κB-responsive firefly luciferase reporter and a constitutively expressed renilla reporter. Blue-light illumination (5 μmol $m^{-2}$ $s^{-1}$) was started 8 h after transfection and samples were fixed for microscopy analysis 24 h later. Representative images are shown; scale bar = 10 μm.

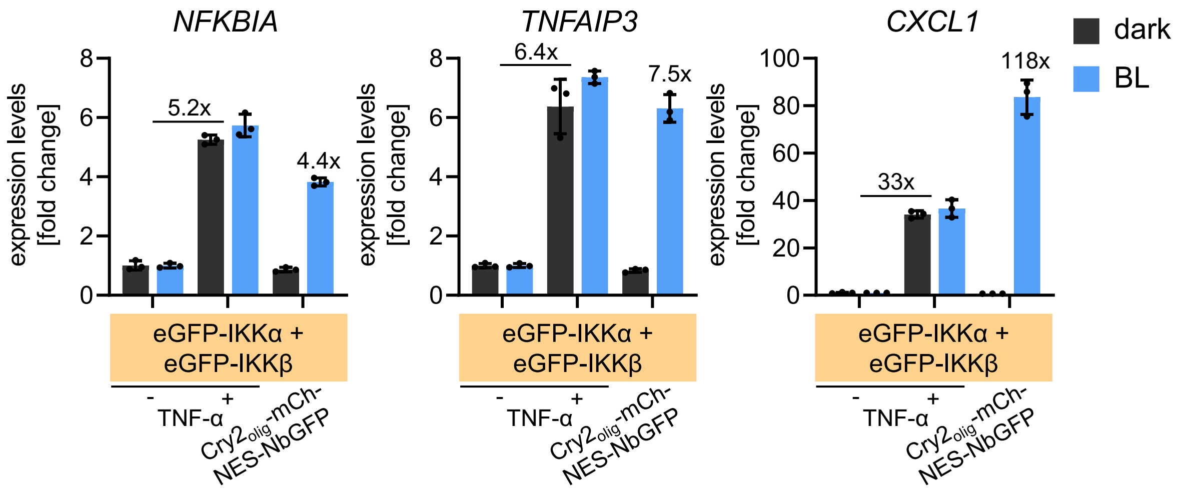


**Supplementary Figure 5: RT-qPCR analysis of optogenetic activation of endogenous NF-κB target genes**

HEK-293T cells were transfected with the indicated constructs. 24 h after transfection, indicated samples were stimulated with 20 ng/ml TNF-α. Simultaneously, blue-light illumination (5 µmol $m^{-2}$ $s^{-1}$) was started. 3 h later, cells were lysed, and total RNA was extracted and reverse transcribed. The expression of three NF-κB target genes was analyzed by qPCR. Data was normalized to the housekeeping gene GUS. Mean $2^{-\Delta\Delta Ct}$ values ± SD are plotted as fold changes to the negative controls (no TNF-α stimulation), N = 3.

**Table S1: Plasmids used in this study**

| Category | Name | Description | Backbone |
|---|---|---|---|
| NbGFP-clustering constructs | pAF352 | Cry2-mCh-NES-NbGFP | pEGFP-C3 |
| | pAF354 | Cry2-mCh-FUS$_N$-NES-NbGFP | pEGFP-C3 |
| | pAF347 | Cry2$_{olig}$-mCh-NES-NbGFP | pEGFP-C3 |
| | pAF300 | Cry2$_{olig}$-mCh-FUS$_N$-NES-NbGFP | pEGFP-C3 |
| | pAF349 | Cry2$_{olig}$-iRFP$_{670}$-NES-NbGFP | pEGFP-C3 |
| | pAF342 | Cry2$_{olig}$-iRFP$_{670}$-FUS$_N$-NES-NbGFP | pEGFP-C3 |
| Constructs to increase clustering | pAF079 | Cry2$_{olig}$-mCh-FUS$_N$-NES | pEGFP-C3 |
| | pAF056 | Cry2$_{olig}$-mCh | pEGFP-C3 |
| | pAF358 | Cry2-mCh-FUS$_N$-NES | pEGFP-C3 |
| NF-κB-POIs | pAF180 | eGFP-IKKα | pEGFP-C3 |
| | pAF181 | eGFP-IKKβ | pEGFP-C3 |
| Reporters | NF-kB Firefly luciferase | 3x NF-κB-RE-Firefly luciferase reporter | Ref. 42 |
| | pAF504 | 3x NF-κB-RE-SEAP reporter | pMF111 |
| | TK-Renilla luciferase | pTK-Renilla luciferase reporter | pRL-TK |
| | CMV-Renilla luciferase | pCMV-Renilla luciferase reporter | pRL-CMV |
| Other | pAF057 | empty vector | pEGFP-C3 |

**Table S2: Transfection conditions of each experiment**

| Figure | Format | Condition | Plasmidname and DNA amount |
|---|---|---|---|
| Figure 1, S2B, S4 | 24-well | eGFP-IKKα + eGFP-IKKβ -/+ TNF-α | pAF180 (6 ng) + pAF181 (6 ng) + NF-κB Firefly luciferase reporter (150 ng) + TK-Renilla luciferase (120 ng) + pAF057 (468 ng) |
| | | eGFP-IKKα + eGFP-IKKβ + NbGFP-clustering construct | pAF180 (6 ng) + pAF181 (6 ng) + pAF352/pAF347/pAF354/pAF300/pAF349/pAF342 (240 ng) + NF-κB Firefly luciferase reporter (150 ng) + TK-Renilla luciferase (120 ng) + pAF057 (228 ng) |
| | | eGFP-IKKα + eGFP-IKKβ + NbGFP-clustering construct + construct to increase clustering | pAF180 (6 ng) + pAF181 (6 ng) + pAF352/pAF347/pAF354/pAF300 (240 ng) + pAF079 (120 ng) + NF-κB Firefly luciferase reporter (150 ng) + TK-Renilla luciferase (120 ng) + pAF057 (228 ng) |
| Figure 2B, C, S1, S2A, S3 | 96-well | -/+ TNF-α | NF-κB Firefly luciferase reporter (25 ng) + CMV-Renilla luciferase (20 ng) + pAF057 (80 ng) |
| | | eGFP-IKKα + eGFP-IKKβ -/+ TNF-α | pAF180 (1 ng) + pAF181 (1 ng) + NF-κB Firefly luciferase reporter (25 ng) + CMV-Renilla luciferase (20 ng) + pAF057 (78 ng) |
| | | eGFP-IKKα + eGFP-IKKβ + NbGFP-clustering construct | pAF180 (1 ng) + pAF181 (1 ng) + pAF352/pAF347/pAF354/pAF300/pAF349/pAF342 (40 ng) + NF-κB Firefly luciferase reporter (25 ng) + CMV-Renilla luciferase (20 ng) + pAF057 (38 ng) |
| | | eGFP-IKKα + eGFP-IKKβ + NbGFP-clustering construct + construct to increase clustering | pAF180 (1 ng) + pAF181 (1 ng) + pAF352/pAF347/pAF354/pAF300/pAF349/pAF342 (40 ng) + pAF079/pAF056/pAF358 (20 ng) + NF-κB Firefly luciferase reporter (25 ng) + CMV-Renilla luciferase (20 ng) + pAF057 (18 ng) |
| Figure 3 | 24-well | eGFP-IKKα + eGFP-IKKβ -/+ TNF-α | pAF180 (6 ng) + pAF181 (6 ng) + pAF504 (150 ng) + pAF057 (588 ng) |
| | | eGFP-IKKα + eGFP-IKKβ + $Cry2_{olig}$-mCh-NES-NbGFP | pAF180 (6 ng) + pAF181 (6 ng) + pAF347 (240 ng) + pAF504 (150 ng) + pAF057 (348 ng) |
